# How long can tardigrades survive in anhydrobiotic state? Searching on tardigrade anhydrobiosis patterns

**DOI:** 10.1101/2022.06.10.495611

**Authors:** Milena Roszkowska, Bartłomiej Gołdyn, Daria Wojciechowska, Zofia Księżkiewicz, Edyta Fiałkowska, Mateusz Pluskota, Hanna Kmita, Łukasz Kaczmarek

## Abstract

Anhydrobiosis is a desiccation tolerance that denotes the ability to survive almost complete dehydration without sustaining damage. However, knowledge on the survival capacity of anhydrobiosis in various species of tardigrades is still very limited. Our research has compared anhydrobiotic capacities of four tardigrade species: *Echiniscus testudo, Paramacrobiotus experimentalis, Pseudohexapodibius degenerans* and *Macrobiotus pseudohufelandi*, which differ in feeding behaviour, habitats occupied, and which belong to different genera. Moreover, in the case of *Ech. testudo* two populations-from urban and natural habitats-were analysed. Clear differences in the anhydrobiotic capacity were observed among different tardigrade species. These differences appear to be determined by the habitat, but not by the nutritional behaviour of the species sharing the habitat type. Moreover, the results indicated that the longer tun state duration, the more time was necessary for the animals to return to activity.

## INTRODUCTION

Cryptobiosis is a latent state of the life cycle in which no signs of an organism’s activity are visible [1]. It is an adaptation to harsh environmental conditions that allow organisms to survive periods unsuitable for active life, e.g., lack of water or very low temperatures. The phenomenon of cryptobiosis has been observed in unicellular (e.g., bacteria and protists), as well as, multicellular (e.g., lichens, liverworts, vascular plants, and some invertebrate animals) organisms. Several different types of cryptobiosis have been described, which are induced in response to specific stress conditions; these include anhydrobiosis (to lack of water), cryobiosis (to low temperature), anoxybiosis (to lack of oxygen) or osmobiosis (change of osmotic conditions) [e.g., 2, 3, 4]. However, the most studied form of cryptobiosis remains anhydrobiosis [e.g., 5, 6, 7, 8, 9, 10].

Anhydrobiosis is defined as dehydration tolerance, which refers to the ability of the species to survive almost complete loss of body water. This adaptation strategy is important in habitats characterized by irregular water availability, like mosses, lichens, liverworts, soil, temporary ponds or puddles, cryoconite holes, or tree holes [e.g., 11, 12, 13, 14]. In multicellular organisms, anhydrobiosis may occur at particular stages of development, but is rarely observed in adulthood [e.g., 2, 3, 4, 6, 8, 15, 16]. Some organisms may stay in an anhydrobiotic state for a very long time [17] and later, may rehydrate and return to active life. It has been suggested that entering anhydrobiosis is a high-cost strategy for an organism, demanding the consumption of extra energy accumulated in specially adapted cells called storage cells [18, 19]. Additionally, previous studies have shown that the longer an organism stays in a dehydrated state, the longer time it requires to return to full activity. Surpassing the dehydration time threshold can lead to lethal outcomes [e.g., 20, 21, 22, 23]. This increased mortality after extended periods of anhydrobiosis may result from an inefficient energy supply and/or insufficiency of the relevant DNA protection/repair mechanisms [10, 16, 18, 19].

Water bears (Tardigrada), roundworms (Nematoda), and wheel animals (Rotifera) are known to be the most effective at entering the anhydrobiosis independently of life stage [e.g., 6, 7, 8, 9, 24, 25] and among these phyla, this process is best studied in tardigrades. In Tardigrada, anhydrobiosis is divided into three stages. The first stage involves dehydration, followed by the second stage, formation of so called “tun”, the fully desiccated animal. The third stage – rehydration-occurs when water becomes accessible for the tardigrade again. Available data on several tardigrade species indicate that the animals differ in their anhydrobiosis capabilities. In general, aquatic species are less efficient in anhydrobiosis than limno-terrestrial ones [e.g., 2, 20, 26]. Since tardigrade tuns may be easily spread by the wind or via vertebrate species such as birds [27, 28], the possibility to colonise new areas exposed to periodic drying [29] may be a good explanation for this difference. However, differences in anhydrobiotic abilities were also observed for species occupying the same habitat [30], but representing different classes, such as Heterotardigrada and Eutardigrada, or with different life strategies [22, 30]. The dehydration conditions and the duration of the tun stage are also crucial parameters underlying the rate and efficiency of return to active life. However, the optimal conditions (like air humidity and temperature) of dehydration and the tun stage duration may differ between species [e.g., 2, 20], as well as, within the same species [e.g., 21, 31, 32].

To date, experiments on anhydrobiosis under controlled conditions have been conducted for several species, including *Acutuncus antarcticus* (Richters, [33]) [34], *Bertolanius volubilis* (Durante Pasa & Maucci, [35]) [36], *Echiniscus testudo* (Doyère, [37]) [38], *Hypsibius exemplaris* Gasiorek, Stec, Morek & Michalczyk, [39] (formerly *Hys. dujardini* (Doyère, [37]) [32, 40, 41, 42, 43, 44, 45, 46, 47, 48]), *Milnesium inceptum* Morek, Suzuki, Schill, Georgiev, Yankova, Marley & Michalczyk, [49] (formerly *Mil. tardigradum* Doyère, [37] [25, 38, 47, 50, 51, 52], *Paramacrobiotus richtersi* (Murray, [53]) [54, 55], *Pam. spatialis* Guidetti, Cesari, Bertolani, Altiero & Rebecchi, [56] [57], *Ramazzottius oberhaeuseri* (Doyère, [37]) [58, 59], *Ram. varieornatus* Bertolani & Kinchin, [60] [40, 43, 61, 62] and *Richtersius coronifer* (Richters, [63]) [58, 59, 64, 65, 66]. However, results of the different experiments cannot be directly compared due to differences in the applied protocols of dehydration, duration of the tun stage and revival monitoring. Moreover, the estimation of anhydrobiosis success was usually limited to calculations of survival rate over a given period of time (most often after one or/and 24 hours), and the animals’ return to activity has generally not been continuously observed. Consequently, interpretation of a general pattern has not been possible.

To obtain comparable results for different species (also those living in different habitats) it is necessary to study anhydrobiosis efficiency in respect to various life strategies under the same standardized and controlled conditions. The only available relevant study conducted under such laboratory conditions was performed by Roszkowska et al. [30] for *Mil. inceptum* and *Ram. subanomalus* (Biserov, [67]), which represent predatory and herbivorous species, respectively, that often co-occur in the same habitat. The authors concluded that the carnivorous species exhibited better anhydrobiosis survivability than the herbivorous one. Their data suggest that within the same habitat, predatory and prey species may adopt different anhydrobiotic strategies. Here, we sought to more closely investigate the anhydrobiosis strategies of tardigrade species that differ in feeding behaviour, habitat preference, and taxonomic status. We conducted a long-term experiment (up to 240 days) to analyse in detail the patterns of tardigrade anhydrobiosis under standardised laboratory conditions. The model organisms were derived from five tardigrade populations (four species) representing Heterotardigrada and Eutardigrada. Since a primary goal of the current study was to assess differences between species inhabiting divergent habitat conditions, we used three species from Poland and one from Madagascar for our experiments. In addition, one species was represented by specimens from two separate populations: an urban population, and a population collected from a national park ca 12 km away from the urban one. Moreover, one of the species is continuously cultured under laboratory conditions. Our results indicate that species differ in their response to dehydration, and further, that their response varies depending on the time spent in anhydrobiosis. Our findings contribute importantly to the mechanistic understanding of anhydrobiosis as a phenomenon with applicative potential.

## MATERIAL AND METHODS

### Species used in experiment and sample processing

Heterotardigrade *Ech. testudo* (populations A and B) and Eutardigrada *Paramacrobiotus experimentalis* Kaczmarek, Mioduchowska, Poprawa & Roszkowska, [68], *Pseudohexapodibius degenerans* (Biserov, [69]) and *Macrobiotus pseudohufelandi* Iharos, [70] were found in moss and soil samples (Table 1). All specimens used in experiments were extracted directly from environmental samples, except of *Pam. experimentalis*, which is cultured under laboratory conditions since 2019 (as described in [71]); those specimens were extracted from the stock culture.

**Table 1.** Species used in experiments and their feeding behaviour and localities.

| Species | Feeding behaviour | Sample type | Locality | Coordinates | Altitude |
| --- | --- | --- | --- | --- | --- |
| *Ech. testudo* (A) | herbivorous | moss on railway embankment | Wielkopolski National Park, Poland | 52°19'09"N;<br>16°48'23"E | 86 m asl |
| *Ech. testudo* (B) | herbivorous | moss on concrete wall | Przybyszewskiego street, Poznań, Poland | 52°24'15"N;<br>16°53'18"E | 87 m asl |
| *Pam. experimentalis* | predatory | moss on soil | Toamasina Province, near Ambavaniasy, Madagascar | 18°56'37"S;<br>48°30'52"E | 717 m asl |
| *Psh. degenerans* | herbivorous | soil between grass roots | Słowiński National Park, Poland | 54°44'54.91"N;<br>17° 26'14.83"E | 8 m asl |
| *Mac. pseudohufelandi* | herbivorous | soil between grass roots | Morasko University Campus, Poznań, Poland | 52°28'04.70"N;<br>16° 55'45.47"E | 88 m asl |

Moss samples were placed into plastic beakers containing 250 ml of tap water. After 18 hours, the water-saturated moss was strongly shaken with tweezers and all plant particles were removed. Water with tardigrades was then poured into a 250 ml plastic cylinder, and allowed to settle for 30 minutes, after which the upper portion of water (ca. 200 ml) was decanted and discarded and the remaining 50 ml was poured into Petri dishes, for tardigrade extraction under a stereomicroscope (Olympus SZ). Soil samples with grass were placed into plastic beakers containing 1000 ml of tap water. After 6-8 hours, the water-saturated soil and grass were strongly shaken with tweezers and then water with floating organic particles was immediately poured through a limnological net with 40 μm mesh size (leaving the grains of sand at the bottom of the beaker). Organic material remaining on the mesh was rinsed with water into a 500 ml beaker and then poured into two 250 ml plastic cylinders. After 30 minutes, the upper portion of water (ca. 200 ml) from each cylinder was decanted and discarded and the remaining 50 ml was poured into Petri dishes, for tardigrade extraction under a stereomicroscope (Olympus SZ).

Specimens were later sorted and only fully active, adult specimens of similar, moderate body length were selected for the anhydrobiosis experiment. Genus abbreviations follow Perry et al. [72].

### Anhydrobiosis experiment

All experiments were performed in 35 mm diameterplastic vented Petri dishes (“experimental dishes”, hereafter) lined at the bottom with white filter paper. Each experimental dish was filled with 450 μl of distilled water and selected individuals (extracted from the environmental samples or culture) were transferred to the experimental dishes using an automatic pipette. Each dish contained seven specimens, and 10 dishes in total were established per species. The experimental dishes were then closed and placed in an environmental chamber with controlled conditions of 40-50% humidity and 20 °C (PolLab, Q-Cell 140, https://www.pol-lab.eu/en/) and allowed to dry over the course of 72 hours. Later, the dishes were stored in the same chamber until the day of rehydration. The drying process took 72 hours (it was necessary for tardigrades to form tuns of correct appearance) and that point was considered as the beginning of the experiment. Specimens were rehydrated after seven different timepoints of tun stage duration (0-, 7-, 14-, 30-, 60-, 120-, 240-days) each constituting a separate experimental group. In the time “0” group, specimens were rehydrated immediately after dehydration (i.e. 72 hours after transferring tardigrades to experimental dishes), whereas specimens from e.g., experimental group “7” were rehydrated after 7 days of tun stage duration, etc.

Rehydration was achieved by adding 3 ml of distilled water to each experimental Petri dish (Fig. 1). Then, specimens were transferred using an automatic pipette to small glass cubes in which they were observed for 10 hours (at room temperature 24 °C) under a stereomicroscope SZ51 or SZX7. After 10 hours, the cubes with non-moving and/or not fully active specimens were placed in an environmental chamber (40-50% humidity and 20 °C) overnight. The next day, they were observed every 30 minutes until 24 hours following rehydration.

**Figure 1.**
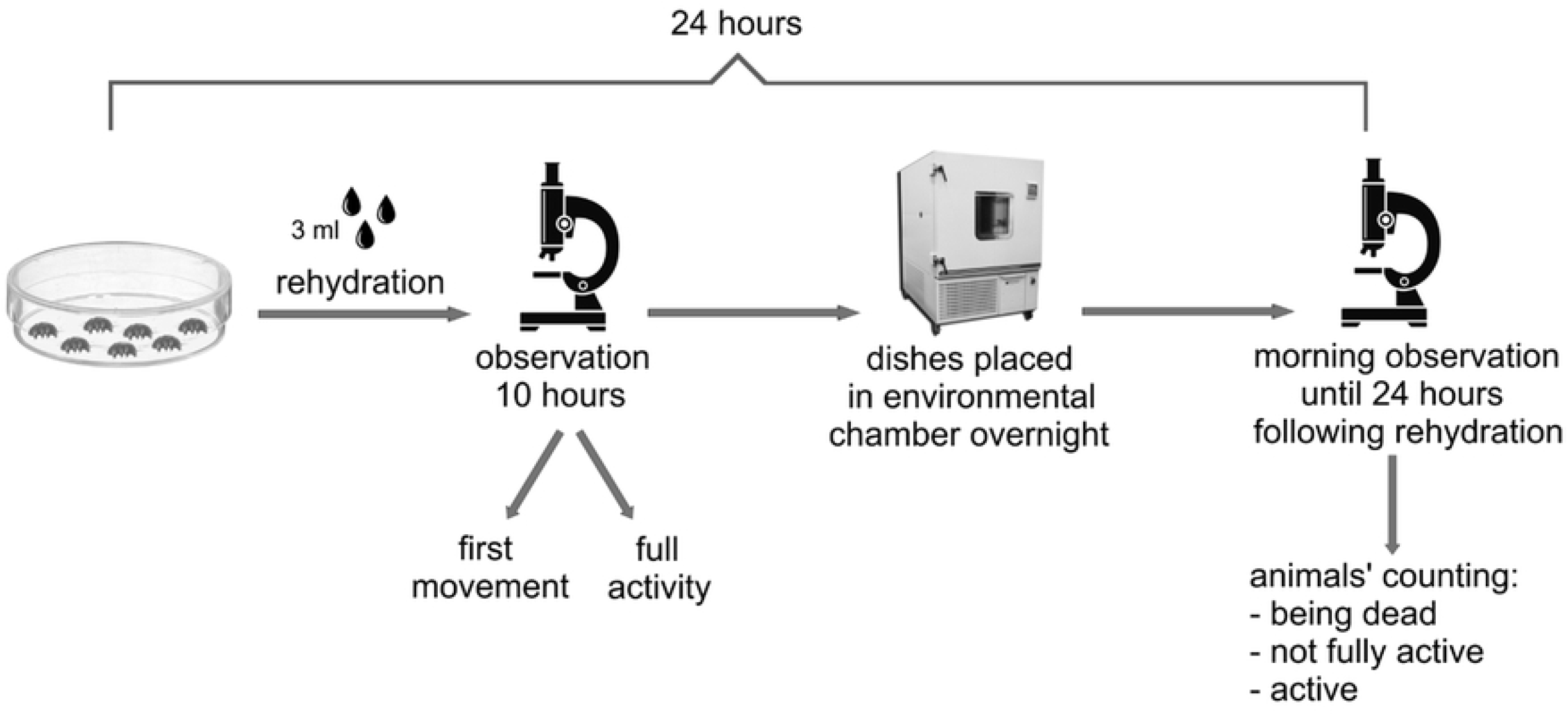
A simple schematic illustration of the experiment (according to Roszkowska et al. [30]).

During observation, the time of both first movement and return to full activity were noted. The first movement was defined as any visible sign of movement of claws, legs, body, buccal tube, etc. Full activity was defined as coordinated movements of the body and legs (i.e., the start of crawling). Specimens observed to be fully active were removed from cubes and no longer observed. All observations were completed after 24 hours and all not fully active specimens or those without any signs of movement (non-moving, interpreted as being dead) were counted.

As described above, seven experimental groups differing in tun stage duration were prepared for each tardigrade species/population. Each experimental group consisted of 10 replicates of 7 specimens each were scored for each time point. Thus, in total, we used 70 specimens per experimental group, for each of the 7 experimental groups, for each tardigrade species/population (i.e., a total of 490 individuals of each species/population).

Seven parameters of survival and activity were measured and used to compare experimental groups and species/populations: (i) SA – success of anhydrobiosis (number and percentage of undamaged specimens, which formed correct tuns, and which survived anhydrobiosis) (ii) the number of non-moving (dead) tardigrades (NM); (iii) the number of individuals that did not reach full activity until the end of the observation (“not fully active”; NFA); (iv) the time required for the first movement of any first individual (FM); (v) the time required for the first movement of all individuals (FMA); (vi) the time required for return to full activity of the first specimen (FA), and (vii) the time required for return to full activity of all individuals (FAA) for a given experimental dish.

### Statistical analysis

Since the distributions of all the recorded variables were far from normality we used nonparametric tests to analyse all the data. Each species/population was tested for the influence of time spent in anhydrobiosis (tun stage duration) on the values for each of six survival and activity parameters described above (NM, NFA, FM, FMA, FA, FAA). Comparisons between experimental groups was done using Permutational Analysis of Variance (Permanova, Anderson 2001) with 999 permutations. Pairwise.adonis test for multiple comparisons adjusted with Bonferroni correction was applied post hoc, to compare differences between individual pairs of experimental groups. The same test was later performed to compare differences in activity measures between the species/populations, after additional stratification for experimental groups. To compare overall differences in activity measurements, a multivariate Permanova was also performed (overall test). All calculations were performed in R 4.0.2 (R Core Team 2020) under RStudio 1.3.1056 using ‘vegan’ package and graphs were produced using ggplot2 package [73]. We considered *p* = 0.05 as the threshold determining statistical significance. Raw data used for all calculations are presented in Appendix S1.

## RESULTS

### General characteristic of anhydrobiosis patterns

The most basic measurement in our studies is survival rate, since ultimately, anhydrobiosis is induced to improve survival. Survival rates are given in Table 2. The highest survival rates (SA) were observed in groups with shorter tun stage durations, from 0–60 days. Exceptions were *Mac. pseudohufelandi* and *Pam. experimentalis*, which exhibited quite high survival rate even after 120 days, at 97% and 86%, respectively. In all species, the lowest survival rates were observed after 240 days of tun stage (1–6% for *Ech. testudo*, 1% for *Psh. degenerans*), though *Mac. pseudohufelandi* and *Pam. experimentalis* exhibited higher 240-day survival rates at 38% and 43%, respectively.

**Table 2.** Values of applied measures of survival and activity used to estimate anhydrobiosis success.

| *Echiniscus testudo* (A) |  |  |  |  |  |  |  |
| --- | --- | --- | --- | --- | --- | --- | --- |
| TR \ MA | NM | NFA | SA | FM (min.) | FMA(min.) | FA(min.) | FAA(min.) |
| 0 | 2 | 7 | 67 (100%) | 3 | 287 | 3 | 287 |
| 7 | 16 | 23 | 54 (93%) | 3 | 155 | 7 | 256 |
| 14 | 10 | 15 | 60 (88%) | 5 | 87 | 7 | 1440 |
| 30 | 8 | 18 | 62 (90%) | 4 | 145 | 9 | 1386 |
| 60 | 11 | 18 | 59 (87%) | 5 | 467 | 9 | 380 |
| 120 | 13 | 21 | 57 (84%) | 14 | 197 | 25 | 301 |
| 240 | 66 | 68 | 2 (6%) | 58 | 67 | 0 | 0 |
| *Echiniscus testudo* (B) |  |  |  |  |  |  |  |
| TR \ MA | NM | NFA | SA | FM | FMA | FA | FAA |
| 0 | 4 | 7 | 65 (94%) | 3 | 68 | 4 | 1244 |
| 7 | 10 | 14 | 60 (88%) | 10 | 86 | 16 | 233 |
| 14 | 6 | 11 | 63 (93%) | 6 | 262 | 9 | 149 |
| 30 | 15 | 27 | 55 (89%) | 10 | 213 | 17 | 1350 |
| 60 | 5 | 7 | 65 (94%) | 5 | 23 | 10 | 390 |
| 120 | 19 | 35 | 51 (76%) | 27 | 231 | 47 | 1440 |
| 240 | 69 | 70 | 1 (1%) | 96 | 96 | 0 | 0 |

*Pseudhexapodibius degenerans*
| TR \ MA | NM | NFA | SA | FM | FMA | FA | FAA |
| --- | --- | --- | --- | --- | --- | --- | --- |
| 0 | 2 | 7 | 68 (99%) | 6 | 1370 | 44 | 1440 |
| 7 | 17 | 21 | 51 (91%) | 8 | 197 | 42 | 380 |
| 14 | 6 | 10 | 62 (95%) | 5 | 222 | 15 | 1423 |
| 30 | 10 | 18 | 58 (97%) | 7 | 402 | 42 | 1440 |
| 60 | 5 | 6 | 64 (93%) | 12 | 1180 | 34 | 1422 |
| 120 | 35 | 52 | 32 (58%) | 21 | 1027 | 101 | 1176 |
| 240 | 69 | 70 | 1 (1%) | 63 | 63 | 0 | 0 |

*Macrobiotus pseudohufelandi*
| TR \ MA | NM | NFA | SA | FM | FMA | FA | FAA |
| --- | --- | --- | --- | --- | --- | --- | --- |
| 0 | 1 | 3 | 66 (99%) | 7 | 221 | 60 | 1343 |
| 7 | 2 | 5 | 67 (100%) | 10 | 312 | 39 | 1451 |
| 14 | 7 | 16 | 61 (90%) | 10 | 1083 | 30 | 1431 |
| 30 | 4 | 21 | 65 (94%) | 12 | 1445 | 31 | 1565 |
| 60 | 7 | 9 | 62 (95%) | 7 | 456 | 22 | 1440 |
| 120 | 2 | 7 | 68 (97%) | 16 | 554 | 61 | 1043 |
| 240 | 43 | 60 | 26 (38%) | 33 | 628 | 313 | 1352 |

*Paramacrobiotus experimentalis*
| TR \ MA | NM | NFA | SA | FM | FMA | FA | FAA |
| --- | --- | --- | --- | --- | --- | --- | --- |
| 0 | 13 | 24 | 55 (100%) | 5 | 1440 | 18 | 1413 |
| 7 | 9 | 16 | 60 (88%) | 7 | 207 | 16 | 1440 |
| 14 | 13 | 32 | 53 (83%) | 10 | 1440 | 62 | 1440 |
| 30 | 10 | 17 | 59 (87%) | 13 | 1440 | 43 | 1440 |
| 60 | 23 | 52 | 46 (67%) | 10 | 573 | 69 | 289 |
| 120 | 12 | 31 | 55 (86%) | 15 | 1359 | 44 | 1332 |
| 240 | 39 | 55 | 29 (43%) | 66 | 1440 | 452 | 1440 |

TR – treatment; MA – measures of activity; SA – success of anhydrobiosis (number of specimens and % of all found, not damaged and forming correct tun specimens); NM – number of non-moving (dead) tardigrades; NFA – number of individuals that didn't reach full activity during time of observation; FM – time required for the first movement of any first individual; FMA – time required for first movement of all individuals; FA – time required for return to full activity of the first individual; FAA – time required for all individuals to return to full activity. Values given in table include all experimental replicate dishes within a given research group for a given duration of the tun stage.

In case of time needed to observe the first movement of any specimen (FM) was quite short in all species/populations for shorter tun stage timepoints (0–60 days, longer after 120 days, and the longest after 240 days of tun stage. The “fastest” species were *Ech. testudo* (A) specimens, which needed only ca. 3 to 5 min to show first movements. After 120 days, all species clearly needed more time. The fastest was *Ech. testudo* (A) which moved at 14 minutes while the slowest *Ech. Testudo* (B) took 27 minutes for FM. The clearest differences in this parameter were visible after 240 days, when species needed between 33 min (*Mac. pseudohufelandi*) and 96 min (*Ech. testudo* (B)).

In some species, full activity of any specimen (FA) was observed after a very short time (*Ech. testudo*, 3–4 min. after the shortest tun stage duration). In contrast, others species required significantly longer time to FA, after even short tun stage periods, such as *Mac. pseudohufelandi* which needed several dozen minutes. Strikingly, after the longest duration of tun stage measured, some species had zero individuals that returned to full activity (i.e., *Ech. testudo* and *Psh. degenerans*), while for others, the time until return was very long (*Mac. pseudohufelandi* – 313 min, and *Pam. experimentalis* – 452 min).

We also measured the time of first movements and full activity of all species in each group, which allows us to assess what a typical or standard behaviour for that species may be. Time of first movements and full activity of all specimens (FMA and FAA, respectively) were rather variable and ranged from several dozen to over a thousand minutes, and thus, were generally not correlated with the duration of the tun stage (Table 2).

### Response to anhydrobiosis of two populations of heterotardigrade Ech. testudo

One outstanding question is whether two different populations of the same species would behave differently. In our study, we sought to address this question by including and comparing two populations of *Ech. testudo*. For most parameters measured, we observed only modest differences between the Wielkopolski National Park (population A) and the Poznań urban population (population B) of *Ech. testudo*. Non-moving (NM) individuals in both groups were very similar in numbers after 0-120 days of tun stage duration, and also at 240 days, when the numbers dropped significantly (Table 2). In natural population A, only two individuals showed any signs of movement after 240 days spent in tun stage, however they did not reach full activity. Similarly, in urban population B, only one individual showed any signs of life after 240 days and it also did not return to full activity (Table 2). Similar numbers of individuals from the two populations failed to return to full activity (NFA), and no significant differences were observed between experimental groups except in the 240 days experimental group. However, in the urban population (population B), the 120-day group had a significantly higher number of NFA specimens than observed for experimental groups of shorter durations of the tun stage (Fig. 2).

**Figure 2.**
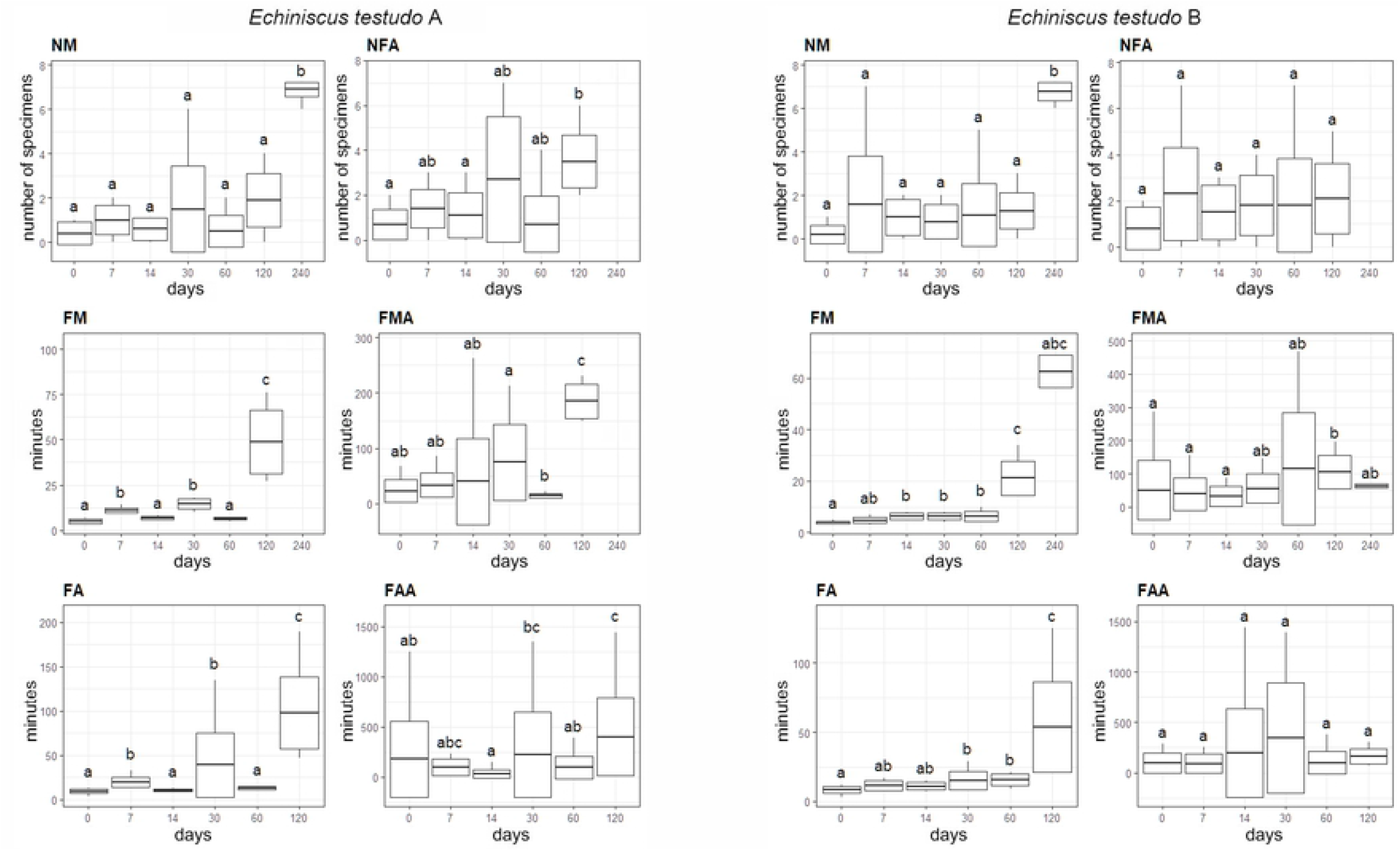
Differences in the number of nonmoving (NM) and not fully active (NFA) individuals, timing of first movement of any individual (FM) and all moving individuals (FMA), full activity of any individual (FA) and full activity of all active individuals (FAA) between particular treatments (horiziontal axis - days in anhydrobiosis) in two populations of *Echiniscus testudo*: A - urban habitat, B - population from surroundings of Wielkopolski National Park.

Times until first movement of any individual (FM) were very similar in both populations: there were no significant differences observed for experimental groups from 0-60 days. However, a difference in FM was evident in the 120 days group and was even stronger for 240 days group. In contrast, despite significant differences between some experimental groups (especially in the urban population), first movement of all individuals (FMA) was a far less informative parameter and did not reveal any differences between populations A and B (Fig. 2).

We found that the relative times to reach full activity for first individual and all individuals (FA and FAA, respectively) were similar to what we observed for FM and FMA. In the 120 days experimental group, FA time was clearly longer than in shorter experimental groups which exhibited only marginal differences between them; none of observed individuals reached full activity after 240 days in the tun stage (Fig. 2). No clear trend in the timing of FAA was observed for either population.

### Response for anhydrobiosis of three eutardigrade species

General patterns in survival and activity measurements in the three eutardigrade species were very similar to patterns described for the two populations of *Ech. testudo*. Animals subjected to the longest duration of the tun stage (240 days) clearly differed from other experimental groups with respect to NM, NFA, FM and FMA, and no clear patterns were observed in the case of FMA and FAA (Fig. 2 and 3). However, in *Pam. experimentalis*, there was no significant difference between the 60 and 240 days tun stage groups in NA and NFA (Fig.3). In *Psh. degenerans*, there was no significant difference in NFA between 120- and 240-days groups, and the intra-group variability in all activity measures for the 30 days experimental group was very high. In the case of *Pam. experimentalis* and *Mac. Pseudohufelandi* significant differences in FA were not observed for between 0-60 days experimental groups and 120 days experimental group but occurred between these groups and 240 days experimental group (Fig. 2 and 3). It should be remembered that this measure analysis was not possible for other species (see above).

**Figure 3.**
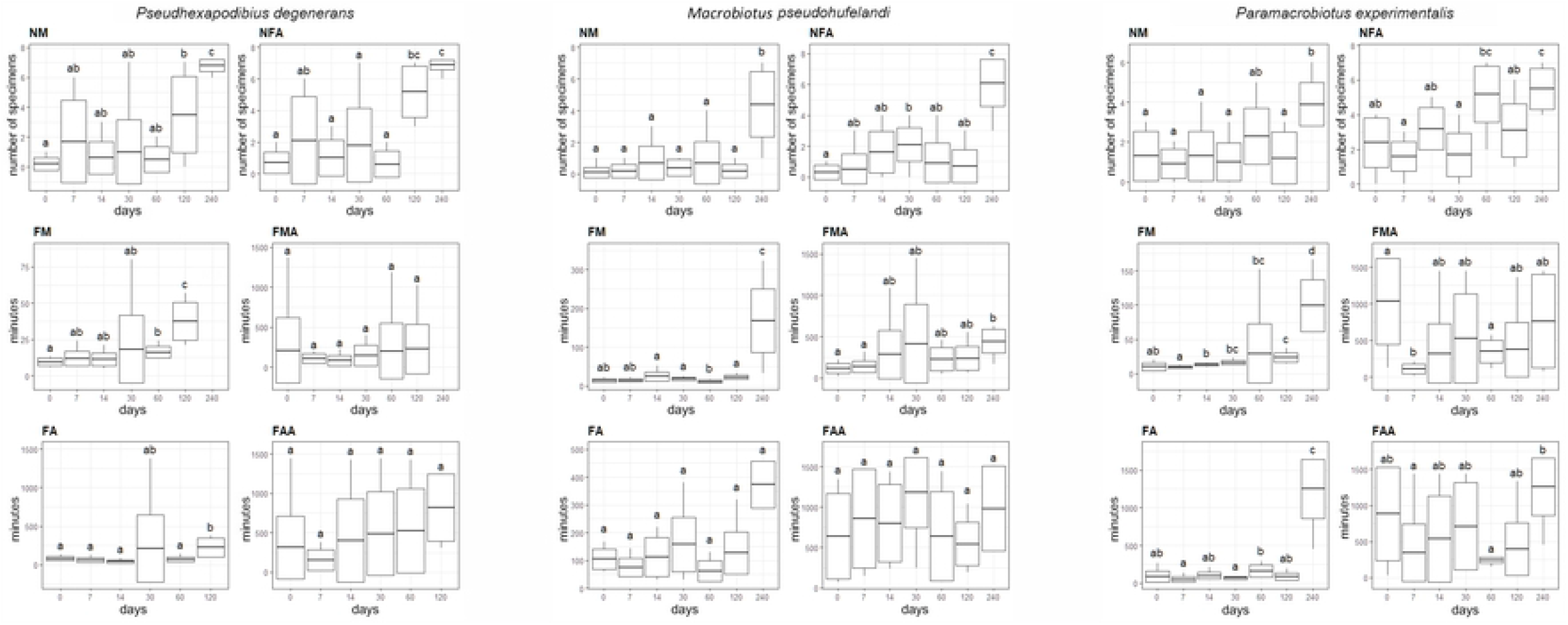
Differences in the number of nonmoving (NM) and not fully active (NFA) individuals, timing of first movement of any individual (FM) and all moving individuals (FMA), full activity of any individual (FA) and full activity of all active individuals (FAA) between particular treatments (horiziontal axis - days in anhydrobiosis) in populations of three tardigrade species: *Pseudhexapodibius degenerans, Macrobiotus pseudokufelandi, Pararamacrobiotus experimentalis*.

### Differences between studied populations

The tests’ values for differences between the five studied populations belonging to the four tardigrade species (Table 3) were statistically significant for all the measures parameters of anhydrobiosis (in all analyses p < 0.001). These differences were the greatest in the case of FA, with F_4,293_ = 109.494, R^2^ = 0.415 and the lowest (but still highly significant) in NFA: F_4,351_ = 7.751, R^2^ = 0.077.

**Table 3.** Differences in the applied activity measures and the measures combined between pairs of populations.

| Species | *Ech. testudo* (A) | *Psh. degenerans* | *Mac. pseudohufelandi* | *Ech. testudo* (B) | *Pam. experimentalis* |
| --- | --- | --- | --- | --- | --- |
| *Ech. testudo* (A) | - | 0.002 | 0.001 | 0.945 | 0.168 |
| *Psh. degenerans* | 6.554 | - | 0.078 | 0.008 | 0.001 |
| *Mac. pseudohufelandi* | 11.829 | 2.334 | - | 0.001 | 0.001 |
| *Ech. testudo* (B) | 0.040 | 5.607 | 10.685 | - | 0.174 |
| *Pam. experimentalis* | 1.455 | 12.633 | 21.152 | 1.666 | - |

NFA
| Species | *Ech. testudo* (A) | *Psh. degenerans* | *Mac. pseudohufelandi* | *Ech. testudo* (B) | *Pam. experimentalis* |
| --- | --- | --- | --- | --- | --- |
| *Ech. testudo* (A) | - | 0.41 | 0.002 | 0.417 | 0.002 |
| *Psh. degenerans* | 3.069 | - | 0.15 | 0.252 | 0.001 |
| *Mac. pseudohufelandi* | 6.626 | 1.37 | - | 0.033 | 0.001 |
| *Ech. testudo* (B) | 0.588 | 0.986 | 3.359 | - | 0.001 |
| *Pam. experimentalis* | 5.868 | 13.697 | 23.223 | 8.825 | - |

FM
| Species | *Ech. testudo* (A) | *Psh. degenerans* | *Mac. pseudohufelandi* | *Ech. testudo* (B) | *Pam. experimentalis* |
| --- | --- | --- | --- | --- | --- |
| *Ech. testudo* (A) | - | 0.001 | 0.001 | 0.001 | 0.001 |
| *Psh. degenerans* | 27.506 | - | 0.087 | 0.001 | 0.639 |
| *Mac. pseudohufelandi* | 44.87 | 3.859 | - | 0.001 | 0.215 |
| *Ech. testudo* (B) | 7.340 | 4.939 | 13.617 | - | 0.001 |
| *Pam. experimentalis* | 35.406 | 1.731 | 0.596 | 8.91 | - |

FMA
| Species | *Ech. testudo* (A) | *Psh. degenerans* | *Mac. pseudohufelandi* | *Ech. testudo* (B) | *Pam. experimentalis* |
| --- | --- | --- | --- | --- | --- |
| *Ech. testudo* (A) | - | 0.001 | 0.001 | 0.375 | 0.001 |
| *Psh. degenerans* | 17.547 | - | 0.001 | 0.001 | 0.001 |
| *Mac. pseudohufelandi* | 51.832 | 18.392 | - | 0.001 | 0.005 |
| *Ech. testudo* (B) | 0.665 | 23.92 | 58.313 | - | 0.001 |
| *Pam. experimentalis* | 47.304 | 23.89 | 5.278 | 51.363 | - |

FA
| Species | *Ech. testudo* (A) | *Psh. degenerans* | *Mac. pseudohufelandi* | *Ech. testudo* (B) | *Pam. experimentalis* |
| --- | --- | --- | --- | --- | --- |
| *Ech. testudo* (A) | - | 0.001 | 0.001 | 0.001 | 0.001 |
| *Psh. degenerans* | 103.43 | - | 0.007 | 0.001 | 0.512 |
| *Mac. pseudohufelandi* | 135.29 | 6.022 | - | 0.001 | 0.164 |
| *Ech. testudo* (B) | 3.037 | 56.388 | 79.949 | - | 0.001 |
| *Pam. experimentalis* | 89.897 | 51.363 | 2.067 | 53.467 | - |

FAA
| Species | *Ech. testudo* (A) | *Psh. degenerans* | *Mac. pseudohufelandi* | *Ech. testudo* (B) | *Pam. experimentalis* |
| --- | --- | --- | --- | --- | --- |
| *Ech. testudo* (A) | - | 0.001 | 0.001 | 0.244 | 0.001 |
| *Psh. degenerans* | 21.247 | - | 0.001 | 0.001 | 0.15 |
| *Mac. pseudohufelandi* | 68.979 | 18.905 | - | 0.001 | 0.012 |
| *Ech. testudo* (B) | 1.0939 | 21.499 | 60.085 | - | 0.001 |
| *Pam. experimentalis* | 33.76 | 2.960 | 6.447 | 31.486 | - |

**All parameters combined (overall test)**
| **Species** | *Ech. testudo* (A) | *Psh. degenerans* | *Mac. pseudohufelandi* | *Ech. testudo* (B) | *Pam. experimentalis* |
| --- | --- | --- | --- | --- | --- |
| *Ech. testudo* (A) | - | 0.001 | 0.001 | 0.21 | 0.001 |
| *Psh. degenerans* | 23.859 | - | 0.001 | 0.001 | 0.001 |
| *Mac. pseudohufelandi* | 66.806 | 16.144 | - | 0.001 | 0.002 |
| *Ech. testudo* (B) | 1.066 | 22.448 | 60.023 | - | 0.001 |
| *Pam. experimentalis* | 46.115 | 10.12 | 5.574 | 42.257 | - |

Upper diagonal: *p* values; Lower diagonal: F values; *NM*—number of non-moving (dead) tardigrades; *NFA*—number of individuals that didn't reach full activity during time of observation; *FM*—time required for the first movement of any first individual; *FMA*—first movement of all individuals; *FA*—time required for return to full activity of the first individual; *FAA*—time required for return to full activity of all individuals.

We found that correlations between species/populations and time spent in the tun stage (experimental groups) was significant across almost all analysed parameters (p < 0.001, F > 1.995, R^2^ > 0.119), with the exception of NM, which did not reach statistical significance (p = 0.062, F_24,315_ = 1.4198, R^2^ = 0.08538). These results indicate that across all remaining parameters, not only did the populations differ in response to drying, but also the patterns of this response changed differently depending on time spent in the tun stage.

Interestingly, the two studied populations of *Ech. testudo* did not differ significantly in their overall anhydrobiosis ability (multivariate model with all measures stratified for time: p = 0.21, F_1,116_ = 1.066, R^2^ = 0.009). However, significant differences between populations were observed in comparing FM (p < 0.001, F_1,121_ = 7.340, R^2^ = 0.058) and FA (p < 0.001, F_1,116_ = 3.037, R^2^ = 0.026).

In all remaining pairs of tardigrade populations, differences in overall anhydrobiosis ability were highly significantly different (p < 0.001, F > 5.574, R^2^ > 0.043) (Table 3). Differences were most pronounced in *Ech. testudo* (A) vs *Psh. degenerans* and *Ech. testudo* (A) vs *Mac. pseudohufelandi* pairs, which differed strongly across all measured parameters. The smallest differences were observed between *Psh. degenerans* and *Pam. experimentalis*, comparing NM, NFA and FMA parameters. FMA was the strongest discriminant of anhydrobiosis response across all species, differing significantly between all the species pairs compared (see Table 3).

## DISCUSSION

Most published studies to date that have reported on the anhydrobiotic abilities of tardigrade species have focused on their ability to enter anhydrobiosis for a short time (from a few hours to 1-3 days). These studies primarily focused on testing survival of tardigrades in environmental extremes (e.g., radiation, changes in humidity, or high temperatures) or repeated anhydrobiosis at different life stages, and included research on DNA repair mechanisms and stress proteins synthesis also due to genomic analyses [21, 25, 32, 34, 36, 40, 41, 43, 44, 45, 46, 47, 51, 52, 54, 57, 61, 65, 66, 74]. The conclusion from these studies is that different tardigrade species have a high degree of survival after short periods of anhydrobiosis, oscillating around 80–90%.

Experiments involving slighting longer periods in tun stage (i.e. 7 days) were conducted by Hengherr et al. [50] for *Milnesium inceptum* (former *Mil. tardigradum*). These specimens were able to survive dozens of cycles of rehydration from the tun stage followed by dehydration for 7 days. When compared with a hydrated control, the periodically dried animals showed a similar longevity as the control. This study showed that the time spent in the tun stage is not “counted” by the animals and lends support to the “Sleeping Beauty” hypothesis [50]. A 7 day tun stage was also applied in experiments designed to test thermotolerance of anhydrobiotic *Ram. varieornatus* [62]. Control groups not exposed to high temperatures had a survival rate of 96–99%, however survivability declined drastically for the tuns kept at temperatures greater than 55 °C. Another 7 day tun stage experiment was conducted on *Hys. exemplaris*, showing differences in survivability between naturally and artificially dried specimens. However, the survivability of studied specimens overall was rather low and oscillated between few to 50% [48].

A 12 day tun stage was tested on Italian and Swedish populations of *Ram. oberhauseri* and *Ric. coronifer*, where similar survival rates were observed for the same species from different locations (∼40% for *Ric. coronifer* and ∼66% for *Ram. oberhaeuseri*). Another study showed that body size had a strong effect on the possibility of returning from anhydrobiosis, but the effect had an opposite direction in case of studied species [58], and this latter study was later confirmed by Faurby et al. [59], except for some differences in survival rates between *Ram. oberhaeuseri* from Sweden and Italian populations and no such differences were found for *Ric. coronifer*. The authors suggested that a geographic variation in successful anhydrobiosis may be a general feature. Furthermore, studies based on a 12 days tun stage showed that medium-sized tardigrades exhibiting better energetic condition enjoy higher survival rates than larger specimens [64].

Experiments addressing the influence of hypomagnetic conditions on *Ech. testudo, Hys. exemplaris* and *Mil inceptum* undergoing anhydrobiosis showed a very high survival rate after 21 days in anhydrobiosis for *Ech. testudo* and *Mil inceptum* (>70%) and very low for *Hys. exemplaris* (20–26%) in control groups not exposed to hypmagnetic conditions. Specimens exposed to hypomagnetic conditions had survival rates of 54–68% for *Ech. testudo* and *Mil inceptum*, and 2-20% for *Hys. exemplaris* [38, 42]. Similarly low survival rates were showed for *Hys. exemplaris* in 7-days experiments by Poprawa et al. [48], who also suggested that it is probably a naturally occurring situation for this freshwater species. In contrast to *Hys. exemplaris*, a very high survival rate (∼ 90%) was reported for *Mil. inceptum*, after 30 and 60 days in anhydrobiosis [47].

Another category of studies focused on tardigrade anhydrobiosis are experiments conducted in outer space. Anhydrobiotic *Ric. coronifer, Ram. oberhauseri* and *Ech. testudo* exposed to space vacuum for two weeks were unambiguously able to return to active life. However after two years in a dehydrated state, none of the tardigrades exposed to cosmic radiation returned to life [75]. Similarly to the laboratory experiments involving high doses of radiation and low temperatures, where tardigrades were desiccated in moss substrate together with rotifers and nematodes, but the survival rate in this lab experiment was rather low [75]. Moreover, Rebecchi et al. [55] tested anhydrobiotic *Pam. richtersi* during TARSE (Tardigrade Resistance to Space Effects), measuring their survival rate after two weeks. They found that survival was very high and differed depending on whether animals were dehydrated in leaf litter (78.9%) or on a paper (94.4%). However, in these experiments the drying protocols for the two comparators were not identical and were not standardized for all species, and the anhydrobiosis time was rather short.

Much longer and confirmed successful anhydrobiosis in tardigrades was reported in a few other papers. In a long-term, semi-natural experiment, tardigrades were desiccated in lichen samples and stored under ambient laboratory conditions. The lichen sample was checked 20 times over 1604 days (i.e., 4 years and 5 months) and the survival rate was calculated for eutardigrade *Ram. oberhaeuseri* and two heterotardigrade species *Ech. trisetosus* [76] and *Ech. testudo* [22]. During this time, *Ram. oberhaeuseri* and *Echiniscus* spp. experienced decreased survivability from 91% and 72% at the beginning of experiment, respectively, to almost 0% at its end. Thus, specimens of *Ram. oberhaeuseri* survived up to 1604 days, while *Echiniscus* spp. up to 1085 days. Baumann [77] reported a successful anhydrobiosis of a *Macrobiotus* [78] species after almost 7 years in the tun stage. Much later, Guidetti and Jönsson [79] analysed eggs of *Ramazzottius* [80] from 9-year-old dried moss and lichen samples which hatched successfully and survived for up to 40 days. Roszkowska et al. [17] reported successful survival of a tun stage lasting 12 and 15 years for *Mac*. cf. *hufelandi* and *Mil. argentinum* Roszkowska, Ostrowska & Kaczmarek, [81], respectively. However, the longest and best-documented survival from the tun stage was reported for heterotardigrade *Ech. testudo*, which was stored dehydrated for ca. 20 years [82]. Unfortunately, most of these experiments (except [22]) were somewhat anecdotal observations of single specimens and were not conducted under strict laboratory conditions. They involved stored dried moss or lichen containing tardigrades, and the experiments consisted merely of checking whether any individuals would return to activity after an extended period in a dry state. Therefore, it is not possible to infer from them any patterns of anhydrobiosis.

Here, we present results of a long-term anhydrobiosis laboratory experiment for four tardigrade species and analyse their ability to recover from dehydration using survival and activity measures of anhydrobiosis success. The parameters measured include: survival of anhydrobiosis (SA), the number of dead (non-moving; NM) specimens as well as measures of animal activity after rehydration, i.e. the number of not fully active (NFA) individuals, the time that individuals required to display the first movement (FM, FMA) and to reach full activity (FA, FAA). The experiments were performed simultaneously for two populations of heterotardigrade *Ech. testudo* (an herbivorous species from moss) and three eutardigrade species, i.e., *Pam. experimentalis* (a predatory species from moss), *Psh. degenerans* (an herbivorous species from soil) and *Mac. pseudohufelandi* (an herbivorous species from soil). Species/populations were collected mostly in Poland, from xerothermic habitats, except for the *Pam. experimentalis* sample, which was collected in a tropical forest in Madagascar and is cultured under laboratory conditions (as described in [71]).

In general, we observed differences in survival rate between species. In studied species the highest survival rate (more than 80% and without visible decreasing) was observed for 0–60 days in the tun stage for *Ech. testudo* population B and *Psh. degenerans* and for 0–120 days in the tun stage for *Ech. testudo* population A, *Mac. pseudohufelandi* and *Pam. experimentalis*. This is in agreement with the very low number of non-moving/dead (NM) specimens for anhydrobiosis times from 0 to 120 days. The lack of decrease in the survival rate with the increasing duration of of the tun stage was also observed for *Pam. richtersi* [31]. However, in that experiment naturally desiccated tardigrades were rehydrated after up to 21 days. Even more spectacular results were obtained for *Mil. inceptum* and *Ram. subanomalus* which exhibited no significant change in survival rate even after 240 days in the tun stage [30]. Strikingly, we observed a drastic decrease in survival after 240 days in the tun stage for all species examined. More specifically, only a few percent of specimens from the *Ech. testudo* population and *Psh. degenerans* and ∼40% of *Mac. pseudohufelandi* and *Pam. Experimentalis* survived after a 240-day tun period.

Interspecific variation in tun stage survival among tardigrades is well known (e.g., [59]). The proposed reasons include body size and/or energetic conditions; in general, larger species/specimens have been found to tolerate anhydrobiosis better and with higher rates of their survival, but the opposite effect has also been observed [58], [64]. Our results seem to support the link between increased survival and increased specimen size, since *Mil. inceptum, Pam. experimentalis, Ram. subanomalus* and *Mac. pseudohufelandi*, which are considered medium sized/larger species exhibited high or very high survival rate after 240 days in the tun stage, as compared to either *Ech. testudo* population (very low survival rate after 240 days in the tun stage). However, it should be also mentioned that *Psh. degenerans* is considered a medium-sized species but had a very low survival rate in our experiment after 240 days in the tun stage. Moreover, *Ech. testudo* belongs to Heterotardigrada and this also may be an explanation of the lower survival rate in comparison to other species which belong to Eutardigrada. The other possible explanation for the variation in anhydrobiosis survival is the energy required to enter, sustain, and leave anhydrobiosis [64]. However, this hypothesis could not be addressed with the experiments presented in the current study.

It is worth noting that the low survival rate we observed for *Ech. testudo* in the current study was surprising, given that specimens of this species were previously shown to return to activity after ∼3 years [22], and even after ∼20 years [82] spent in the tun stage. This is rather unexpected and hard to explain in comparison of our results, but it is not possible to compare the success rate in natural samples and our laboratory experiment (semi-natural drying in moss samples by Rebecchi et al. [22] and Jørgensen et al. [82] *versus* artificial drying on filter paper in the present study). Besides differences in the applied protocols, the differences may be caused by genetic factors (although Jørgensen et al. [82] showed low genetic variability in *Ech. testudo* in large geographic range) or phenotypic plasticity of the species in response to microclimatic conditions.

The time needed for the first movement of any specimen (FM) for all studied species, especially for short durations of the tun stage, is very short (maximum few minutes), but much longer after 120 days and especially long after 240 days in the tun stage. The fastest species was *Ech. testudo* specimens which needed only ca. 3 to 10 min., then the other of the species which needed ca. 6 to 13 minutes. Return to full activity of the first specimen was observed also very fast (after shortest time of anhydrobiosis) in some species (e.g., for *Ech. testudo* 3–4 min.) while other species needed much more time (e.g., even several dozen minutes for *Mac. pseudohufelandi*). After the longest time spend in the tun stage (240 days) none of the specimens of *Ech. testudo* and *Psh. degenerans* returned to full activity during 24h of observation. For other species the time of the return was very long 5–7h (*Mac. pseudohufelandi* and *Pam. experimentalis*).

Differences between the five populations of the four tardigrade species were statistically significant for all of the activity measures. Interaction between the species and time spent in the tun stage indicate that in all the remaining measures, not only the populations differed in values of their response to drying, but also the patterns of this response changed differently depending on the time spent in the tun stage. Another interesting observation was that the two populations of *Ech. testudo* in our study did not differ significantly when their overall reaction to anhydrobiosis was compared, however significant differences in the timing of FM and FA in this pair of populations were detected. This is in agreement with the results of Faurby et al. [59] suggesting that populations of the same species from different localities may differ in anhydrobiotic abilities. We suggest that such differences may, over longer time scales, lead to speciation.

It is clearly visible that the longer lasts the tun stage the longer time is required to return to full activity, although based on the present study and the previous publication by Roszkowska et al. [30], it can be concluded that this relation is far from linear one. This is in agreement with explanations that time spent in the tun stage is correlated with the number of damages which need to be repaired as well as the time required to activate metabolism [see e.g., 23, 55, 57]. Accordingly, the time required to reach full activity after shorter durations of the tun stage, for all species/populations, is relatively short (within the range of minutes). Such a rapid response in nature may represent an adaptation to the brief periods of liquid water occurrence in tardigrades’ natural habitats. Moss cushions, lichens, or soil itself (at least on the surface) may absorb water in the early morning when fog and dew appear. When the mosses, lichens and topsoil subsequently dry out when the sun rises, there is a rapid decline in air and substrate humidity. Thus, the liquid water phase in these habitats can last only a few hours per day or even less, with longer hydroperiods only occurring occasionally under rainy conditions. However, even after rains, especially in temporary or dry climatic zones, water is frequently available for only a few hours or a few days. Thus, rapid recovery from anhydrobiosis can be regarded as an adaptation to this type of temporary habitats since — in this short amount of time — tardigrades must supplement energy resources exploited during anhydrobiosis (e.g., [83]) and carry out their entire life cycle. This could also suggest that returning to active life faster than other tardigrades would be a favourable trait, since fast-recovering animals could begin feeding and reproducing earlier, exploiting potentially limited local resources and helping ensure greater reproductive success.

Our study also noted that a predatory species (*Pam. experimentalis*) needed more time to return to activity after tun stage than herbivorous species (like *Ech. testudo*), which contradicts observations made by Roszkowska et al [30] which were explained by a prey-predator strategy. However, it needs to be stated that, in contrast to the research by Roszkowska et al [30], in present studies herbivorous and predator species were not collected from the same sample and belong to the different genera.

In conclusion, we showed clear differences in the anhydrobiotic abilities between different species/populations reflected by the observed different patterns. The patterns appear to be determined by the habitat, but not in the case of the same species population and by the way of nutrition for species sharing the habitat. Consistent with previous reports, we found that a critical factor is the duration of the tun stage, whose duration generally in inversely correlated with the general success of and time required for recovery from anhydrobiosis. The underlying mechanisms of this tendency are still not fully understood, but they seem to be directly related to ecological strategy supported by metabolism.

## ETHICS APPROVAL AND CONSENT TO PARTICIPATE

Not applicable.

## CONSENT FOR PUBLICATION

Not applicable.

## AVAILABILITY OF DATA AND MATERIALS

Data generated and analyzed during this study are included in this published article.

## AUTHORS’ CONTRIBUTIONS

MR, ŁK and HK came up with research ideas. HK and ŁK supervised the performed analyses. MR, BG, DW, ZK, EF, MP, HK and ŁK wrote the final version of the manuscript. MR, DW, ZK, EF and ŁK carried out the tardigrade cultures and collected animals for experiments. MR, DW, ZK and ŁK performed experiments with anhydrobiosis. MR, MP and BG prepared microphotographs and figures. MP and BG prepared statistical analyses. All authors read and approved the final manuscript.

## ACKNOWLEDGEMENTS

The work was supported by the research grant of National Science Centre, Poland, NCN 2016/21/B/NZ4/00131. Studies have been conducted in the framework of activities of BARg (Biodiversity and Astrobiology Research group). The authors also wish to thank Cambridge Proofreading LLC (http://proofreading.org/) for their linguistic assistance.

## SUPPORTING INFORMATION

Appendix S1. The raw data used for all statistical calculations.

